# Distractor suppression does and does not depend on pre-distractor alpha-band activity

**DOI:** 10.1101/2023.07.18.549512

**Authors:** Zach V. Redding, Ian C. Fiebelkorn

**Affiliations:** Department of Neuroscience and Del Monte Institute for Neuroscience, University of Rochester, Rochester, NY 14627, USA

## Abstract

Selective attention enhances behaviorally important information and suppresses distracting information. Research on the neural basis of selective attention has largely focused on sensory enhancement, with less focus on sensory suppression. Enhancement and suppression can operate through a push-pull relationship that arises from competitive interactions among neural populations. There has been considerable debate, however, regarding (i) whether suppression can also operate independent of enhancement and (ii) whether neural processes associated with the voluntary deployment of suppression can occur prior to distractor onset. We provide further behavioral and electrophysiological evidence of independent suppression at cued distractor locations while humans performed a visual search task. We specifically utilize two established EEG markers of suppression: alpha power (∼8–15 Hz) and the distractor positivity (P_D_). Increased alpha power has been linked with attenuated sensory processing, while the P_D_—a component of event-related potentials—has been linked with successful distractor suppression. The present results demonstrate that cueing the location of an upcoming distractor speeded responding and led to an earlier onset P_D_, consistent with earlier suppression due to strategic use of a spatial cue. We further demonstrate that higher pre-distractor alpha power contralateral to distractors was generally associated with successful suppression on both cued and non-cued trials. However, there was no consistent change in alpha power associated with the spatial cue, meaning cueing effects on behavioral and neural measures occurred independent of alpha-related gating of sensory processing. These findings reveal the importance of pre-distractor neural processes for subsequent distractor suppression.

**Significance Statement:** Selective suppression of distracting information is important for survival, contributing to preferential processing of behaviorally important information. Does foreknowledge of an upcoming distractor’s location help with suppression? Here, we recorded EEG while subjects performed a target detection task with cues that indicated the location of upcoming distractors. Behavioral and electrophysiological results revealed that foreknowledge of a distractor’s location speeded suppression, thereby facilitating target detection. The results further revealed a significant relationship between pre-stimulus alpha-band activity and successful suppression; however, pre-stimulus alpha-band activity was not consistently lateralized relative to the spatially informative cues. The present findings therefore demonstrate that target detection can benefit from foreknowledge of distractor location in a process that is independent of alpha-related gating of sensory processing.

## Introduction

The brain uses selective attention to preferentially process behaviorally important aspects of the environment^1,2^. This preferential processing occurs through both enhancement and suppression. The enhancement of behaviorally important sensory information is based on a combination of stimulus salience (i.e., stimulus properties), previous experience, and behavioral goals (i.e., voluntary control)^3–7^. In comparison, there is still considerable debate regarding the variables and neural mechanisms that contribute to the suppression of distracting information^8–13^. This debate largely focuses on a few interrelated topics, including (i) whether suppression can occur independent of neural processes associated with enhancement^3,9,12–14^, (ii) whether suppression can be guided by previous experience and/or voluntary control^15,16–18^, and (iii) whether suppression can occur in anticipation of a predictable distractor^8,19^. Here, we used spatially informative cues during a visual search task to investigate voluntary suppression of distractors, independent of enhancement.

There is growing evidence that neural processes occurring prior to the onset of predictable distractors can contribute to successful suppression^20–25^, but there are still disagreements whether such anticipatory suppression is under voluntary control. Noonan et al.^20^, for example, demonstrated that presenting a distractor at a consistently cued location led to both faster target detection and electrophysiological evidence of distractor suppression (i.e., a lower amplitude P1 component in the ERP). Distractor cueing, however, was not associated with distractor suppression when the location of the spatial cue was varied from trial to trial. The authors therefore concluded that distractor suppression could be based on predictive information derived from experience^26–29^ but not on flexible, voluntary mechanisms of cognitive control (like those linked to the enhancement of behaviorally important information^3,14,30,31^).

Contrary to this conclusion, several behavioral studies have shown that an informative spatial cue, presented at different locations on different trials, can promote distractor suppression^22,23,32^. A recent EEG study provided neural evidence that the strategic use of informative cues is linked to attenuated processing of distractors^25^. Knowing the location of an upcoming distractor was associated with (i) higher anticipatory alpha power over visual cortex, contralateral to the cued distractor location, and (ii) a lower-amplitude N2pc in response to the distractor^25^. Higher alpha power has been repeatedly linked to attenuated sensory processing^33–39^, and the N2pc is an ERP component that reflects the covert enhancement of targets^40–43^. These findings therefore provide evidence that the strategic use of spatial cues can reduce attentional capture by a distractor. As a complement to the N2pc component, the P_D_ component reflects the covert suppression of distractors^8,44^. Additional EEG evidence of voluntary suppression has shown that cueing the location of an upcoming distractor is associated with reduced P_D_ amplitude^24,45^, an effect that covaried with relative increases in alpha power contralateral to the spatial cue^45^. The authors interpreted these results as evidence that anticipatory mechanisms (associated with alpha lateralization prior to distractor onset) reduce the need for reactive distractor suppression.

Despite such behavioral and electrophysiological evidence, there remains skepticism regarding voluntary suppression of distractors^9,11–13,46^. Here, we further tested the effects of a spatially informative cue on distractor suppression (as indexed by the P_D_). We also tested whether and how suppression-related neural activity occurring prior to distractor onset (i.e., alpha power lateralization relative to a cued distractor location) affects successful suppression. While there is considerable evidence that increased alpha power is associated with attenuated sensory processing^33–39^, there is mixed evidence for a relationship between increased alpha power and voluntary distractor suppression^13,20,25,27,47–49^. The present results (i) provide evidence of voluntary suppression, occurring independent of enhancement, and (ii) reveal a reliance of successful suppression on neural processing that occurs prior to distractor onset. While higher alpha power contralateral to distractors was generally associated with successful suppression, behavioral and electrophysiological evidence of voluntary suppression occurred without consistent cue-related changes in alpha power. The present results therefore demonstrate that voluntary suppression can be mediated by mechanisms that occur independently from alpha-related gating of sensory processing^33,34^.

## Methods

### Subjects

Twenty-seven individuals with normal or corrected-to-normal vision and no history of neurological disease participated in the experiment (21 females, 6 males; mean age = 25.3 years). All subjects provided informed consent. The study was conducted in accordance with protocols approved by the Research Subjects Review Board at the University of Rochester.

### Apparatus and Stimuli

Subjects were seated in a comfortable chair in a sound and light-attenuated chamber. Chair and table height were adjusted so that the subject could comfortably use padded chin and forehead rests. A foot stool was provided if needed. Subjects were instructed to remain as relaxed as possible during trials to minimize noise in the EEG attributable to muscle activity. Task contingencies were controlled with custom Presentation (Neurobehavioral Systems, Inc.) scripts run on a Dell Precision 5820 desktop computer with a 27-inch Acer Predator XB2 LCD monitor (1920 × 1080 pixels at 240 Hz) positioned 57 cm from subjects’ eyes. Figure 1 depicts the primary experimental display which consisted of a white (187 cd/m^2^) fixation square (0.5° x 0.5°) and a visual search array comprising four stimuli of varying shapes (circle, square, diamond, hexagon) drawn in red (49.5 cd/m^2^, x =.536, y = .342) or green (46.2 cd/m^2^, x =.309, y = .548) on a black background. Stimuli were positioned 4° from the fixation square (above, right, below, left). The circle had a diameter of 1.4°, the square had a width of 1.24°, and the diamond and hexagon had similar areas (all within 10 pixels). Each shape in the search array contained a black line (0.0625° × 0.75°) rotated 45° clockwise or counterclockwise from horizontal.

**Figure 1.**
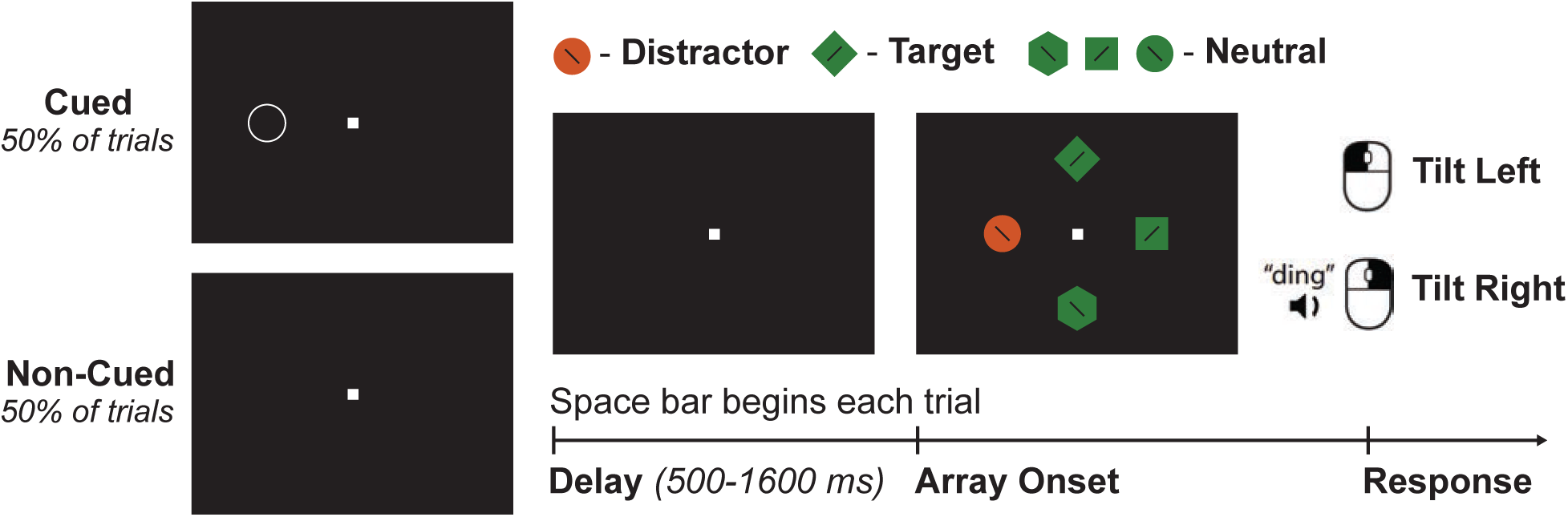
Experimental task. A peripheral cue indicated the location of a distractor that occurred on 50% of trials. The space bar was pressed to initiate a variable duration delay period (500– 1600 ms). The left or right mouse buttons were used to indicate the orientation of a tilted line within a target of fixed color and shape, regardless of the presence or absence of the distractor.

### Procedure and Design

Each trial started with a cueing period in which a tone sounded after 500 ms to signal that subjects could continue the trial when ready. On 50% of trials subjects saw only a white fixation square on a black background (non-cued), while on the remaining 50% of trials a cue was presented at one of the four stimulus locations (i.e., top, right, bottom, left), indicating where a distractor could appear in the search array (cued). On cued trials with a distractor, the distractor always occurred at the cued location (i.e., 100% cue validity). The distractor cue was an unfilled white circle with an outside diameter of 1.4° and an internal diameter of 1.3° (i.e., a line thickness of 0.1°). To isolate the effects of distractor suppression, target location always remained unknown prior to array onset, preventing subjects from relying on anticipatory target enhancement^50^. When ready, subjects advanced the trial by pressing the space bar on a standard computer keyboard. The screen then displayed only the fixation square for a duration sampled randomly from a uniform distribution between 500–1600 ms. Following this delay period, the search array was displayed and remained on screen until subjects responded. Subjects searched the array to find a target shape presented with either three randomly selected neutral shapes of the same color (20% of trials) or two randomly selected neutral shapes of the same color and one randomly selected color-singleton distractor shape of a different color (80% of trials). These conditions were randomized across trials within a block. Previous research has demonstrated different conditions under which a singleton distractor either can or cannot be successfully suppressed, as indexed by the P_D_ component and behavioral performance. Here, we wanted to test the influence of voluntary attentional deployment on the successful suppression of distracting information. We therefore utilized an experimental design that has previously been shown to promote successful suppression of a color singleton. That is, target features and distractor color were held constant and non-target shapes were heterogenous^51^. The target shape was either a circle or a diamond drawn in either red or green. Target shape and color were counterbalanced across subjects. After identifying the target shape within the array, subjects discriminated the orientation of the line inside the target shape and responded by pressing the left or right mouse button. Correct responses triggered a feedback tone. 600 ms after the response, the search array disappeared and the cueing period for the next trial began. After every 20 trials, the error rate for those trials was displayed on the screen. Subjects were instructed to respond as quickly as possible while maintaining error rates of 10% or less. Each subject completed at least 20 practice trials followed by 8 blocks of 140 trials for a total of 1120 trials. Subjects were instructed to maintain fixation without blinking for the duration of each trial (i.e., the period beginning with the subject pressing the space bar and ending with their response). Fixation was monitored using an infrared eye tracking camera (EyeLink 1000 Plus, SR Research Ltd., Ontario, Canada). If fixation was broken, the trial was aborted and a warning message appeared, instructing subjects to press the space bar to begin the next trial. If trials were aborted in this way, additional trials were added to the block such that the total number of completed trials for each condition was consistent across subjects.

### EEG Recording and Preprocessing

EEG data were collected using a BioSemi ActiveTwo system (Biosemi) from 128 active Ag/AgCl electrodes. Data were digitized at 2048 Hz and downsampled to 512 Hz offline. All electrode impedances were kept below 20 kΩ. To ensure consistent placement of the EEG cap, the vertex electrode (A1) was placed at 50% of the distance between the inion and the nasion and between the tragus on the left and the right ears. EEG data were analyzed using the FieldTrip toolbox (Oostenveld et al., 2011) and custom scripts written in MATLAB (R2021b). Offline, the continuous EEG data were re-referenced to the average of all electrodes, high-pass filtered (4^th^ order Butterworth with 0.5 Hz cutoff), then segmented into epochs relative to search array onset (-2.2 to 0.9 s). Bad channels were visually identified. Artifacts related to blinks and eye movements were excluded from the data by design, as task performance was contingent on maintaining fixation (i.e., trials with blinks and saccades were aborted). Data were visually inspected for each subject and channel to determine a voltage threshold for identifying noise transients within the delay period or the ERP window. Bad data were interpolated using a distance-weighted average of the four nearest good electrodes, but only if the average distance of these electrodes was less than 4.0 cm (otherwise data were excluded). On average 2.37% of data were interpolated and kept, while 0.67% of data were excluded (0.22% of data from the primary electrodes of interest, A10/B7, were excluded from ERP analyses).

### Behavioral Analyses

Only trials with RTs less than 2000 ms were included in analyses. For analyses of RT, only correct trials were used. Repeated measures ANOVA was performed on RT and accuracy measures with distractor presence (present/absent) and cue presence (cued/non-cued) as repeated measures factors. To facilitate analyses of suppression-related ERP components (i.e., the P_D_), distractor-present trials outnumbered distractor-absent trials (4:1). For each subject, trials were therefore randomly subsampled without replacement from distractor present conditions to match that subject’s average trial count for distractor absent conditions. Analyses were performed on the subsampled data. For analyses of cueing effects on RT as a function of delay duration, all cued and non-cued trials were used after averaging across distractor present and distractor absent conditions (i.e., there was no subsampling). For each subject, trials were binned by delay duration. Overlapping bins (200 ms width) were created at 50-ms increments (e.g., 500-700 ms, 550-750 ms, …, 1400-1600 ms) and RTs were averaged within these bins. Differences between cued and non-cued trials at each of these bins were measured using a permutation-based approach to compensate for multiple comparisons^52^. Briefly, mean RTs were compared for cued versus non-cued trials within each bin using paired-samples *t*-tests. For each subject, trials were randomly reassigned to cued or non-cued conditions and the process of calculating means and performing *t*-tests for each bin was repeated. The maximum *t-*value (*t*_max_) across all bins was recorded. This process of shuffling the data and recalculating *t*_max_ across bins was repeated 1000 times. The resulting distribution of *t*_max_ values was then used to determine the likelihood of observed *t*-values given the null hypothesis of no difference between conditions.

### ERP Analyses

ERP analyses were limited to the time window from -100 to 400 ms relative to array onset. Time series were baseline corrected using the period from -100 to 0 ms and averaged across trials. For each subject, ERPs were calculated at electrodes A10/B7 (see Figure 3D & 3E), contralateral and ipsilateral to lateralized positions of interest. These electrodes correspond with electrodes PO7/8, which have been typically used to measure the P_D_ component^19,44,48,53,54^. Specifically, ERPs were calculated (i) relative to the position of the singleton distractor on distractor present trials (i.e., cued/distractor present and non-cued/distractor present trials) and (ii) relative to the cued location on cued/distractor absent trials. For each condition, difference waves were calculated for each subject by subtracting the ipsilateral ERP waveform (i.e., relative to the cue/distractor location) from the contralateral ERP waveform. To achieve this, electrode positions were flipped across hemispheres for trials with the cue/distractor on the right side. Data for these trials were then combined with data for trials with the cue/distractor on the left side. Only trials with midline targets (i.e., top or bottom) were analyzed, eliminating any effects of target processing on lateralized ERP components^42^. For plotting, data were low-pass filtered with a cutoff of 30 Hz. The amplitudes of ERP components were calculated using signed area to minimize biases from the choice of measurement windows^52,55^. Following prior work^53^, P_D_ amplitude was determined by calculating the positive area from 100–300 ms and N2pc amplitude was determined by calculating the negative area from 200–400 ms. To test the significance of the observed components, we used a nonparametric permutation approach that was developed by Sawaki and colleagues^55^. For each of 1000 iterations, contralateral and ipsilateral labels were randomized across trials and ERPs were recalculated. The signed areas (positive for P_D_ and negative for N2pc) were calculated within their respective windows for each of these iterations, resulting in a distribution of signed areas that would be expected if the null hypothesis were true (i.e., there was no P_D_ or N2pc component). Statistical significance was then determined using the percentage of permuted results that surpassed the observed signed areas. For comparisons of component amplitudes between conditions (i.e., cued vs. non-cued), paired-samples *t*-tests were performed on the observed signed areas. To test for differences in the component latencies between conditions, 50% fractional area latencies were calculated for each subject^52^, followed by paired-samples *t*-tests.

### Determination of Subject-Specific Peak Alpha Frequency

Alpha frequency varies between substantially individuals^56^; therefore, peak alpha frequencies were determined for each subject. All alpha power analyses used data from electrodes A10/B7, A15/A28, A16/A29, and A17/A30, a cluster that corresponds to electrodes PO3/4, PO7/8, and O1/2, which have been used in prior work studying the role of alpha in attentional selection^48,57^. Data for trials with delays longer than 850 ms were isolated from -500 to 0 ms relative to array onset, and IRASA^58^ was used to obtain oscillatory power spectra at electrodes contralateral to cued locations. Data were padded with zeros to obtain a frequency resolution of 0.1 Hz. For each subject, local maxima in the alpha-band frequency (8–15 Hz) were identified after averaging spectra across trials and electrodes.

### Effects of Pre-Array Alpha Power on ERPs

For each subject, alpha power was measured immediately prior to array onset. EEG data from each trial were convolved with a 2-cycle, complex Morlet wavelet at the subject-specific peak alpha frequency. The temporal envelope of a wavelet is determined by its frequency and width (e.g., a 9-Hz wavelet with 2 cycles is ∼222 ms wide). Observed alpha peak frequencies were greater than 9 Hz for all subjects; therefore, wavelets were centered at -111 ms relative to array onset to avoid overlap with subsequent evoked activity (i.e., the sensory response associated with array onset). Lateralization indices were calculated separately for the four conditions (i.e., cued distractor on the left, cued distractor on the right, non-cued distractor on the left, and non-cued distractor on the right) using the following equation:

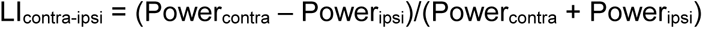

Lateralization indices for each condition were then averaged across electrode pairs, and trials were median split based on the resulting values. ERPs were then separately calculated, using data from electrodes A10/B7, for high and low alpha lateralization trials. Comparisons of component (i.e., P_D_ and N2pc) amplitudes between the high and low alpha lateralization conditions were conducted with paired-samples *t*-tests, using signed area during the same windows used for prior ERP analyses (see above).

### Analyses of Alpha Power Lateralization

For each subject, data were convolved with a 2-cycle complex Morlet wavelet of subject-specific alpha frequency, centered -111 ms relative to array onset. As contralateral and ipsilateral labels could not be assigned for non-cued trials, we calculated an alpha lateralization index using the following equation:

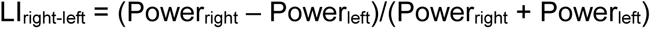

Here, right and left refer to the electrode positions. Lateralization indices were averaged across the subset of electrodes used for analyses of ERP components based on alpha lateralization (see above). For each subject, non-cued trials were subsampled to match the average trial count for cue left and cue right conditions. For each condition, alpha lateralization indices were tested against zero using one-sample t-tests and conditions were compared using one-way ANOVA.

## Results

### Behavioral Evidence of Voluntary Suppression

We first tested whether the presence of the distractor and the cue (see Fig. 1) influenced behavior (i.e., RT and accuracy). For the analysis of RTs, a repeated measures ANOVA revealed no evidence for an interaction between distractor presence and cue presence [F(1,26) = 0.692, p = .413, η_p_^2^ = .026] (Fig. 2A), but there were statistically significant main effects for both distractor presence and cue presence. RTs were faster for distractor-present trials (734 ms) relative to distractor-absent trials (751 ms) [F(1,26) = 11.883, p = 0.002, η_p_^2^ = .314]. Although it may seem counterintuitive that RTs were faster on distractor-present trials, it should be noted that a suppressed distractor decreases the number of stimuli in the array that need to be searched to identify the target^59–62^. That is, on distractor-present trials, subjects only needed to search three locations (rather than four locations), leading to faster RTs. Critically, RTs were also faster for cued trials (732 ms) relative to non-cued trials (753 ms) [F(1,26) = 21.358, p < .0001, η_p_^2^ = .451]. Foreknowledge about the location of an upcoming distractor led to significantly faster RTs, consistent with voluntary suppression of the distractor location.

**Figure 2.**
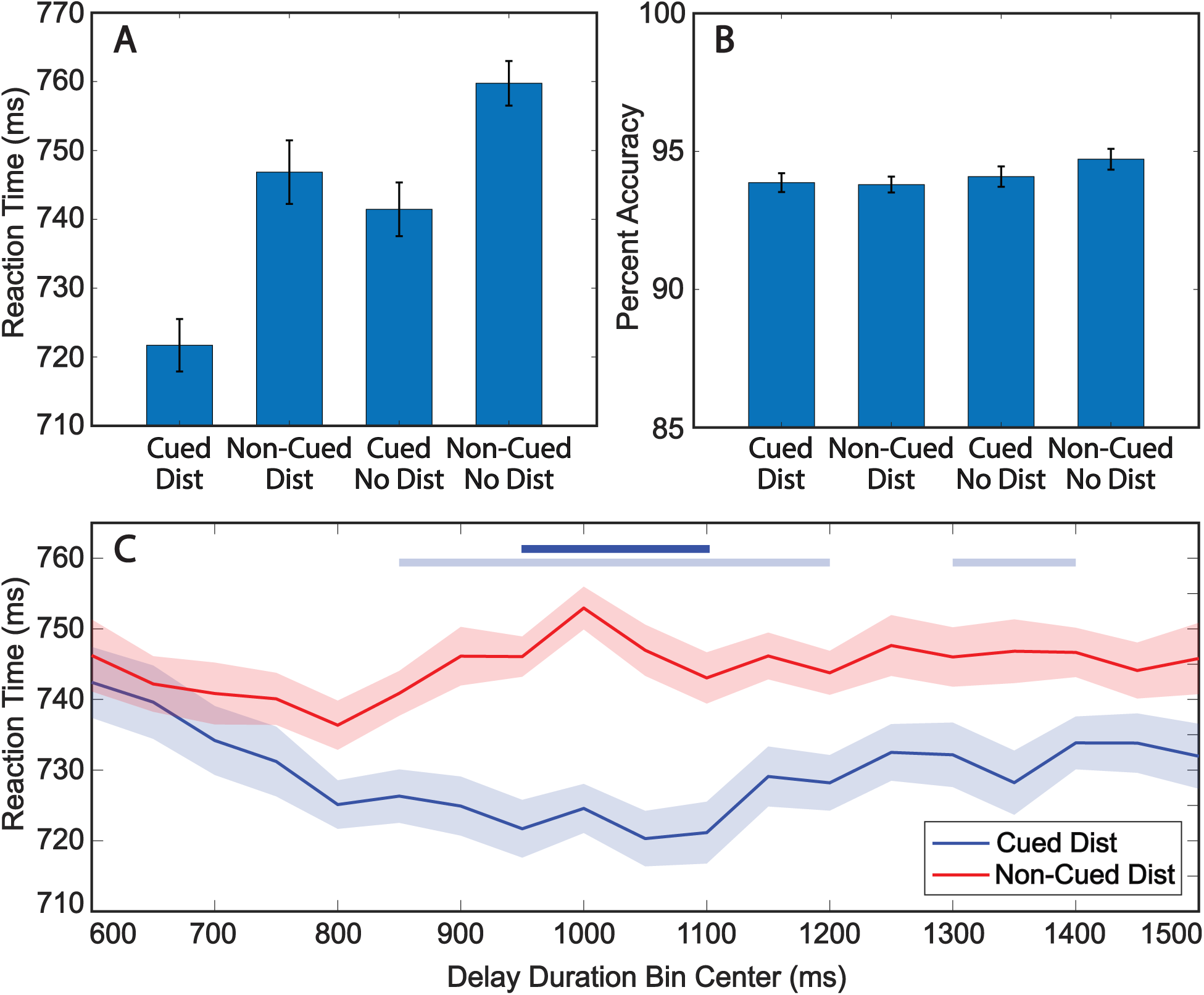
The distractor cue was associated with faster RTs. ***A***, Grand average RTs for combinations of cue (present/absent) and distractor (present/absent) conditions. ***B***, Mean accuracy for combinations of cue and distractor conditions. ***C***, Grand average RTs for cued (blue) and non-cued (red) distractor trials within overlapping 200 ms bins. All error bars indicate ±1 SEM. Bolded light blue lines (above the RT data) indicate the time points at which RTs significantly differed between cued and non-cued trials without correcting for multiple comparisons, while bolded dark blue lines indicate the time points at which RTs significantly differed after correcting for multiple comparisons (p < .05).

For the analysis of accuracy, a repeated measures ANOVA revealed no evidence for an interaction between distractor presence and cue presence [F(1,26) = 1.802, p = 0.191, η_p_^2^ = .065] (Fig. 2B). There was also no evidence for either a main effect of distractor presence [F(1,26) = 0.492, p =0.489, η_p_^2^ = .012] or cue presence [F(1,26) = 1.538, p =0.226, η_p_^2^ = .019] (Fig. 2B). For the remaining analyses, we therefore focused exclusively on RTs.

We next measured the difference between RTs on cued and non-cued trials as a function of delay length (Fig. 2C), characterizing how cueing effects evolved over time. RTs for cued trials were significantly faster (relative to non-cued trials) between 850–1200 ms and 1300–1400 ms after delay onset (p < .05). After corrections for multiple comparisons, differences remained significant between 950–1100 ms. These results suggest that the effects of a spatially informative cue on distractor suppression were not immediate in the present behavioral task, but instead developed after hundreds of milliseconds.

### ERP Evidence of Voluntary Suppression

We next measured the P_D_—an ERP component associated with successful distractor suppression^19,44,53,63^—and determined whether it was different on cued versus non-cued trials. Figure 3A depicts the ERP waveforms contralateral and ipsilateral to cued distractors on the horizontal meridian. Importantly, the target was on the vertical meridian in each of these trials and therefore did not contribute to lateralized ERPs. Figure 3B shows the corresponding ERP waveforms relative to non-cued distractors. Figure 3C shows the contralateral-minus-ipsilateral difference waves for cued and non-cued distractors, isolating the P_D_ component^53^. The area under the difference wave within the P_D_ window was significantly positive for both cued trials (p = .003) and non-cued trials (p < .001). There was no difference in P_D_ amplitude between cued trials (58.0 uVms) and non-cued trials (66.7 uVms) [*t*(26) = 0.688, p = .498, *d* = 0.13]; however, P_D_ onset latency (Fig. 3C) was significantly earlier on cued trials (171 ms) relative to non-cued trials (210 ms) [*t*(26) = 5.833, p < .001, *d* = 1.12]. While subjects were able to successfully suppress the distractor on both cued and non-cued trials (as indexed by the P_D_ component), foreknowledge of the distractor location was associated with earlier distractor suppression, consistent with faster RTs on cued trials (Fig. 2).

**Figure 3.**
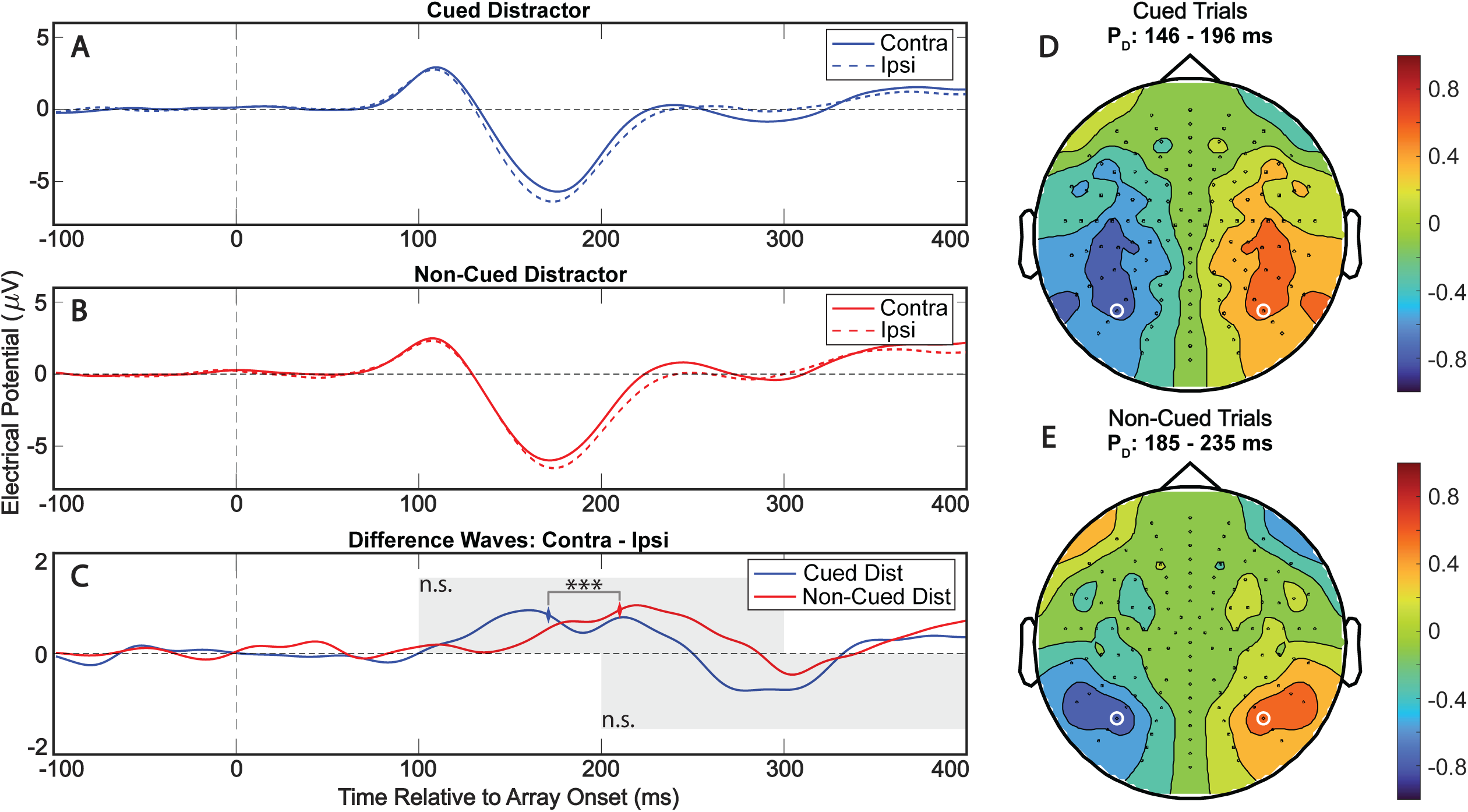
The distractor cue was associated with shorter latency distractor suppression on trials with lateralized distractors and midline targets. ***A***, Grand average ERPs recorded at electrodes contralateral and ipsilateral to lateralized, cued distractors. ***B***, Grand average ERPs relative to lateralized, non-cued distractors. ***C***, Contralateral-minus-ipsilateral difference waves relative to cued (blue) and non-cued (red) distractors. Grey shaded boxes represent analysis windows for P_D_ (100–300 ms) and N2pc (200–400 ms) components. Results of tests comparing the signed area between cued and non-cued conditions are indicated in the corner of the corresponding shaded region (n.s. is not significant). P_D_ latency is indicated by colored diamonds for cued (blue) and non-cued (red) trials. *** p < .001. ***D***, Topographic maps depicting mean amplitude during observed peak P_D_ window for cued trials (146–196 ms). Average voltages for contralateral minus ipsilateral difference wave. ***E***, Topographic maps depicting mean amplitude during observed peak P_D_ window for non-cued trials (185–235 ms).

Figure 4 shows the ERP waveforms contralateral and ipsilateral to targets presented on the horizontal meridian when either cued distractors (Fig. 4A) or non-cued distractors (Fig. 4B) were presented on the vertical meridian. Here, we focused on the N2pc, an ERP component associated with selective enhancement of sensory processing. Figure 4C shows the contralateral-minus-ipsilateral difference waves, isolating the N2pc component. In agreement with previous studies^41,43^, we observed a significant N2pc contralateral to the target. The area under the difference wave within the N2pc window was significantly negative for both cued trials (p = .003) and non-cued trials (p < .001). There was no difference in N2pc amplitude when distractor location was cued (58.0 uVms) relative to non-cued trials (65.4 uVms) [*t*(26) = 0.917, p = .368, *d* = 0.18]. Additionally, N2pc onset latency was not different on cued trials (277 ms) relative to non-cued trials (282 ms) [*t*(26) = 0.601, p = .553, *d* = 0.12]. Combined with ERP analyses of lateralized distractor trials (Fig. 3), these results indicate that cues influenced suppression of the distractor but not target selection (i.e., enhancement).

**Figure 4.**
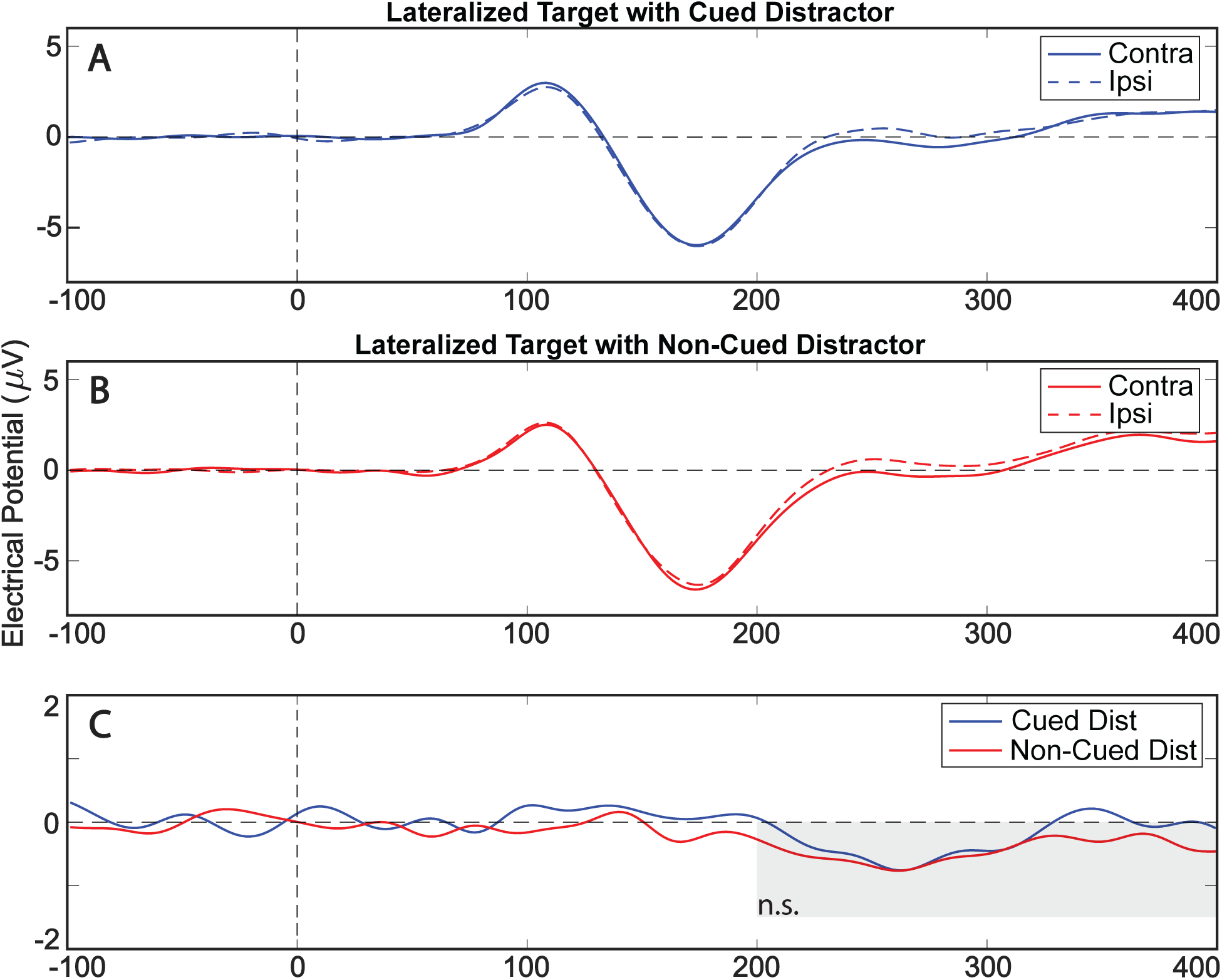
The distractor cue did not lead to differences in target selection on trials with lateralized targets and midline distractors. ***A***, Grand average ERPs recorded at electrodes contralateral and ipsilateral to targets on trials with cued distractors. ***B***, Grand average ERPs recorded at electrodes contralateral and ipsilateral to targets on trials with non-cued distractors. ***C***, Contralateral-minus-ipsilateral difference waves relative to targets on cued trials (blue) and non-cued trials (red). Grey shaded boxes represent analysis windows for the N2pc (200–400 ms) component. Results of the test comparing the signed area between cued and non-cued conditions are indicated in the corner of the shaded region (n.s. is not significant).

A closer look at Figure 3, suggests that the significant P_D,_ associated with distractor suppression, might have been followed by an N2pc-like ERP component, potentially associated with selection of the distractor. ERPs are generated by averaging trials. The P_D_ and the N2pc-like component could therefore be associated with different sets of trials^64^. That is, the P_D_ could have occurred on trials when subjects successfully suppressed the distractor, while the N2pc-like component could have occurred on trials when the distractor instead captured attention (i.e., subjects failed to suppress the distractor). The results of the following analysis provide further evidence for this interpretation. Here, it should be noted, however, the area of this N2pc-like component was not significantly different from zero for cued trials (p = .063) or for non-cued trials (p = .441).

### Alpha power lateralization prior to array onset affects ERP components

Increased alpha power over cortical regions processing behaviorally irrelevant stimuli has been repeatedly associated with sensory suppression^33,35–37,57^. Here, we tested whether the magnitude of alpha lateralization prior to the presentation of the array (i.e., the distractor) was associated with successful distractor suppression. Two of the twenty-seven subjects (7.4%) did not have clear alpha peaks in both the cued and non-cued conditions and were therefore excluded from alpha power analyses. The average alpha peak frequency across subjects and conditions was 11.3 Hz with a SD of 0.9 Hz. Alpha peak frequency was significantly higher in the cued condition (11.4 Hz) relative to the non-cued condition (11.2 Hz) [*t*(24) = 3.508, p = .002, *d* = 0.7]. Subject-specific alpha frequencies for further analyses were therefore based on each subject’s peak frequency for the cued and non-cued conditions, respectively.

Figure 5A depicts ERP waveforms and difference waves for trials with either high or low pre-array alpha power lateralization (i.e., split based on median alpha lateralization) relative to cued distractor locations. P_D_ amplitude was significantly higher on trials with greater alpha activity contralateral to cued distractor locations [*t*(24) = 2.74, p = .011, *d* = 0.55]. In contrast, amplitude of the N2pc-like component was significantly reduced on trials with higher alpha contralateral to cued distractor locations [*t*(24) = 2.513, p = .019, *d* = 0.5]. Figure 5B depicts ERP waveforms and difference waves for trials with high or low pre-array alpha power lateralization relative to non-cued distractor locations. Similar to cued trials, P_D_ amplitude was significantly higher on trials with greater alpha activity contralateral to non-cued distractor locations [*t*(24) = 2.547, p = .018, *d* = 0.51]. Also similar to cued trials, amplitude of the N2pc-like component was significantly reduced on trials with higher alpha contralateral to non-cued distractor locations [*t*(24) = 2.315, p = .03, *d* = 0.46]. These results indicate that higher pre-array alpha lateralization contralateral to the distractor location was associated with successful distractor suppression (as indexed by the P_D_ component). In comparison, lower pre-array alpha lateralization contralateral to the distractor location was associated with attentional capture by the distractor (as indexed by the N2pc-like component). Pre-array alpha lateralization, relative to the distractor location, was therefore an indicator of subsequent distractor suppression. These results are also consistent with the notion that the P_D_ and the N2pc-like component, which are both observable in Figure 3, occurred on different trials. That is, on a given trial, the distractor was either suppressed or selected (not suppressed, then selected).

**Figure 5.**
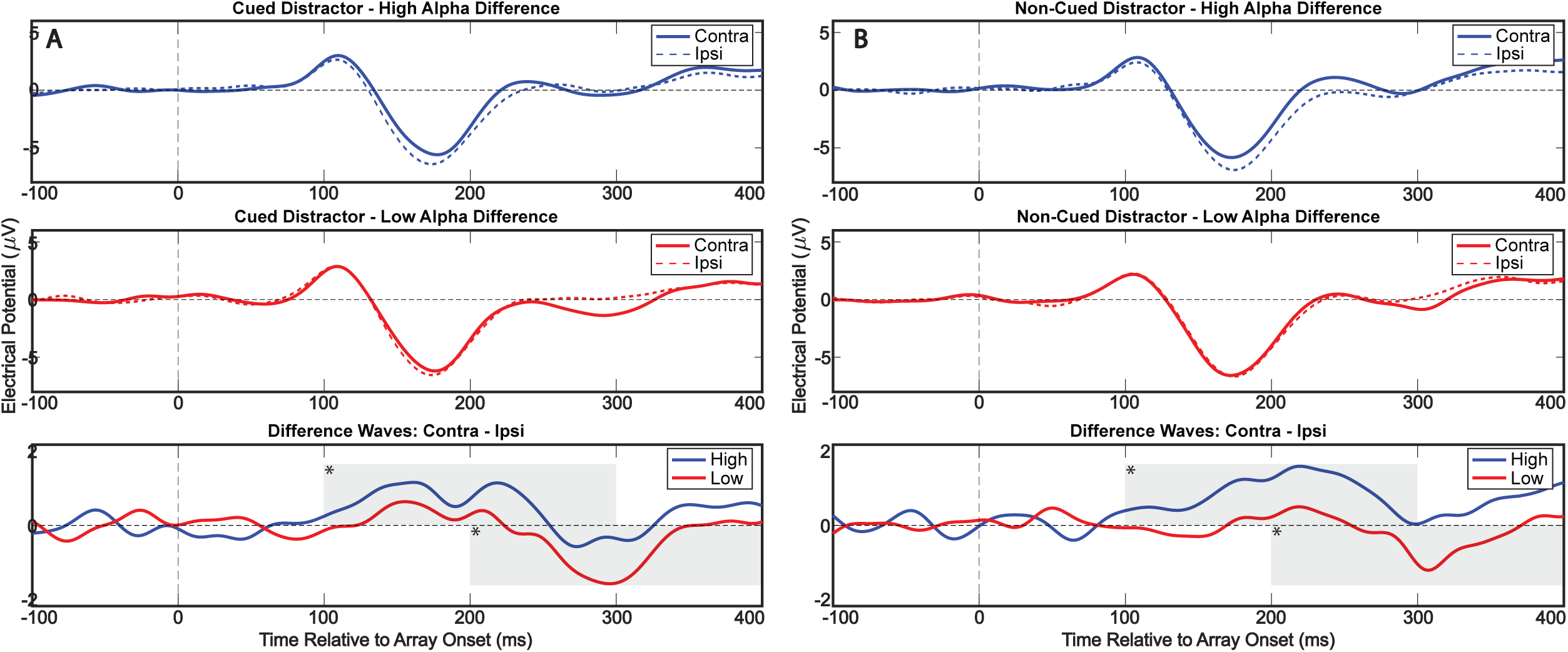
Higher pre-array alpha lateralization was associated with successful distractor suppression. ***A-B***, Grand average ERPs recorded at electrodes contralateral and ipsilateral to cued (***A***) and non-cued (***B***) distractors on trials with higher pre-array alpha power contralateral to the distractor location (top). Grand average ERPs on trials with lower pre-array alpha power (middle). Contralateral-minus-ipsilateral difference waves (bottom). Grey shaded boxes represent analysis windows for P_D_ (100–300 ms) and N2pc (200–400 ms) components. Results of tests comparing the signed area between high and low alpha conditions are indicated in the corner of the corresponding shaded region. * p < .05.

### Pre-array alpha power relative to the cued location

We next tested whether alpha power after the cue and before array onset was consistently higher contralateral to the cued distractor location^25,45^. Figure 6 depicts alpha power lateralization prior to array onset for cue left, cue right, and non-cued trials. Alpha power lateralization was not different from zero for either cue left (LI = -0.011) [t(24) = 0.464, p = 0.647, d = 0.093], cue right (LI = 0.021) [t(24) = 0.958, p = 0.348, d = 0.192], or non-cued trials (LI = -0.0008) [t(24) = 0.037, p = 0.971, d = 0.008]. There was no difference in alpha lateralization between cue left, cue right, and non-cued trials [F(2,48) = 0.419, p = 0.563, η_p_ ^2^ = .017]. These results indicate that the spatially informative cue was not reliably associated with a relative increase in anticipatory (i.e., pre-array) alpha power contralateral to the upcoming distractor location, consistent with other recent studies^27,65^. See below for a detailed discussion of how we interpret these findings.

**Figure 6.**
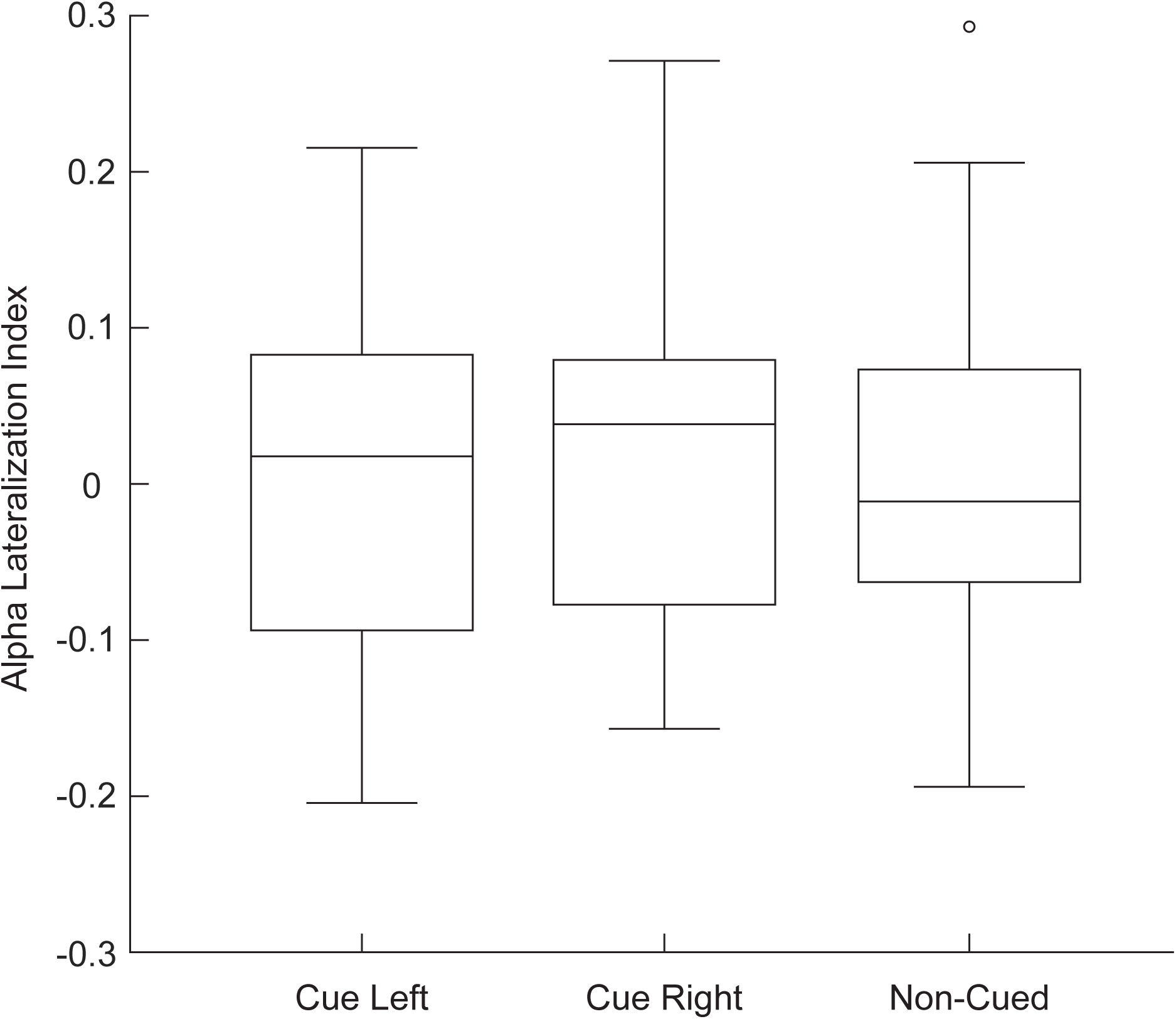
There were no cue-related differences in pre-array alpha power lateralization. The line within each box represents the median, while the upper and lower boundaries of each box represent the upper and lower quartiles, respectively. Lines outside the box represent maximum and minimum data points excluding outliers. Outliers were more than 1.5 times the interquartile range outside of upper and lower quartiles and are denoted with an open circle.

## Discussion

Here, we demonstrated (i) evidence of voluntary suppression, independent of target enhancement, and (ii) evidence that pre-distractor (or anticipatory) neural activity can influence subsequent suppression independent of changes in alpha-band activity. Subjects responded to targets faster when a salient distractor was presented. This counter-intuitive speeding of responses on trials with distractors—reported here and elsewhere^59–62^—has been posited to reflect successful suppression of the distractor location, thereby reducing the number of positions that must be searched to locate a target^8^. Importantly, responses were fastest when the location of a potential distractor was cued in advance, indicating that subjects strategically used the cue to more effectively suppress the distractor. ERP analyses further revealed an earlier onset P_D_ component—a proxy for successful suppression—on cued trials, indicating that the presence of the cue was associated with earlier suppression of the distractor location. Consistent with the involvement of pre-distractor neural activity in successful distractor suppression, we only measured a significant P_D_ component on trials when alpha power immediately prior to array onset was relatively higher contralateral to the distractor location. This link between alpha lateralization and the presence of the P_D_ component, however, was present on both cued and non-cued trials. Relatively higher alpha power contralateral to the subsequent distractor location was therefore more generally associated with successful suppression, while the presence of the spatially informative cue was associated with earlier suppression. We did not observe a consistent increase in alpha-band lateralization associated with the presence of the cue, suggesting that the observed cue-related speeding of suppression in the present task occurred independently of alpha-related gating of sensory processing^33,34^.

Color singleton distractors, like those used here, can either be suppressed or capture attention^59,60^, depending on various factors (e.g., whether the color of the singleton is consistent or variable). The likelihood that a distractor will be successfully suppressed is an important consideration when studying neural mechanisms associated with selective processing^50^. Here, we designed our task to test the influence of voluntary processes on successful suppression (see also^24^). We therefore used a consistent distractor color across trials, which has previously been shown to increase the likelihood of successful suppression^65^. Because of this consistent distractor color, subjects may have suppressed the distractor using not only the voluntary deployment of spatial attention (i.e., based on the spatially informative cue) but also feature-based learning^12^ (i.e., based on the consistent distractor color). While the use of feature-based learning in the present experimental task is speculative, faster responses on cued trials (Fig. 2) indicate that subjects did voluntarily suppress the distractor based on location. The present results therefore add further support to accumulating evidence that suppression can be voluntarily deployed during selective spatial processing, leading to behavioral benefits^22,23,25,32^. It should be pointed out, however, that several studies have reported no behavioral benefits of cueing a distractor location^20,28^, including during a similar visual search task^28^. The factors contributing to whether distractor locations can be voluntarily suppressed include, but are likely not limited to, cue validity^28^, similarities between targets and distractors^66^, prior engagement of attention at the to-be-suppressed location^67–69^, implicit learning of distractor features^22,23,25,32^, the proximity of targets and distractors^25^, the relative salience of target and distractor stimuli^70^, and similarities between targets and other non-target or neutral stimuli^71^.

While several studies have demonstrated behavioral evidence of voluntary suppression^22,23,25,72^, there is relatively little neural evidence that suppression can be voluntarily (and flexibly) deployed on a trial-by-trial basis, independent of processes associated with enhancement^9,12,13^. In a recent EEG study, distractor cues reduced the amplitude of the N2pc component observed in response to salient distractors^25^. The N2pc component reflects attention-related selection, meaning that the spatially informative cue reduced attentional capture. In comparison, the present results demonstrate that a spatially informative cue can decrease the latency of successful suppression, as indexed by the latency of the P_D_ component. Prior studies have demonstrated that cueing a distractor location can reduce the amplitude of the P_D_ component (i.e., on cued trials relative to non-cued trials)^24,45^. Those studies have interpreted this reduced amplitude as evidence that participants were suppressing the distractor location prior to distractor onset. That is, those studies interpret the P_D_ component as reflecting reactive suppression that is no longer necessary if a spatially informative cue enables pre-distractor (or anticipatory) suppression. The present findings show no cue-related change in P_D_ amplitude but a decreased latency (Fig. 3) and faster response times (Fig. 2). Our findings are therefore consistent with the P_D_ component reflecting successful suppression, regardless of whether that suppression involves anticipatory or reactive processes^48,65^.

Alpha-band activity (∼8–15 Hz) seems to play a key role in selective sensory processing, potentially inhibiting task-irrelevant brain regions^33,34,37^ and/or aligning neural activity across brain regions to facilitate communication^34,73–75^. Alpha-band activity can track the deployment of covert spatial attention, with power increasing ipsilateral to attended locations (i.e., contralateral to unattended locations)^33,35–37,57,76,77^. Such findings suggest that increased alpha power is associated with the inhibition of unattended locations to prevent distraction; however, much of the existing evidence to support the role of alpha-band activity in distractor suppression comes from studies in which suppression was conflated with target enhancement. In other words, putative suppression-related changes in alpha-band activity could be an indirect effect of target enhancement^50^. Several studies have now measured changes in alpha-band activity while manipulating distractor suppression orthogonally to target enhancement, demonstrating preparatory changes in alpha power associated with expectation suppression^48^ and voluntary suppression^25,78^. Here, we did not observe consistent increases in alpha power contralateral to the location of the distractor cue, despite observing a more general relationship between alpha lateralization and successful suppression (Fig. 5). These results are consistent with several recent reports in which no consistent alpha power increases were seen contralateral to the location of an expected distractor^27,65^. Further research will be needed to parse the conditions under which a spatially informative cue leads to consistent alpha lateralization.

While we did not observe relatively increased alpha power contralateral to the cued location, we did observe a relationship between pre-distractor alpha lateralization and successful distractor suppression. Specifically, P_D_ amplitude was greater when alpha power was relatively increased contralateral to the distractor location immediately prior to array onset, for both cued and non-cued trials. These results support a strong link between pre-distractor neural activity, specifically alpha-band activity, and subsequent distractor suppression^25,45^. Splitting trials based on pre-array alpha lateralization further dissociated trials that resulted in either successful distractor suppression or distractor-driven attentional capture. As stated previously, the N2pc component is typically associated with attentional selection (rather than suppression); however, other studies have reported an N2pc following the P_D_ component^54,79–84^. This counterintuitive pattern could be explained by the P_D_ and N2pc components occurring on separate trials that are then merged together in the grand-averaged ERP waveform^8^. That is, trials when the distractor was successfully suppressed contribute to the P_D_ component, while trials when the distractor resulted in attentional capture contribute to the N2pc component. The present results are consistent with this interpretation, as P_D_ and N2pc components observed in response to singleton distractors were dissociated after splitting trials by pre-array alpha lateralization. The N2pc component was weaker on trials with a stronger P_D_ component, consistent with successful suppression. In contrast, the N2pc was stronger on trials with a weaker P_D_ component, consistent with attentional capture (i.e., a failure to suppress the distractor).

The present findings provide behavioral and electrophysiological evidence that voluntary, selective processing can contribute to distractor suppression, independent of voluntary, selective processing that contributes to target enhancement. Although we observed a link between pre-distractor alpha power and successful suppression, alpha power was not consistently modulated in response to the spatially informative cue. The cue-related changes in distractor suppression observed in the present task (i.e., faster RTs and a shorter latency P_D_ component) were therefore not mediated through alpha-related mechanisms. The extent to which suppression-related changes in alpha-band activity are tied to neural processes associated with enhancement remains a topic of debate^47^. In the present task, we observed an absence of cue-related enhancement (as indexed by the N2pc component) and an absence of cue-related changes in alpha lateralization. It is unknown, however, whether their combined absence is related. Regardless, the present findings provide evidence that voluntary suppression, independent of both target enhancement and alpha-related sensory gating, contributes to selective attention, helping to determine which aspects of the environment will receive preferential processing.

## Acknowledgments

This work was supported by grants from the National Science Foundation (NSF 2120539), the National Institutes of Health (NEI R01EY033726), and the Searle Scholars Program to I.C.F.

## Notes

**Conflict of interest:** The authors declare no competing financial interests.

### Competing Interest Statement

The authors have declared no competing interest.

